# Multi-modal microRNA and non-targeted proteome analysis of liquid biopsies from the distal lung collected by Particles in Exhaled Air (PExA) reveal presence from extracellular vesicles

**DOI:** 10.1101/2024.11.12.620657

**Authors:** M. Bonatti, M. Ezerskyte, P. Lindberg, B. Zöhrer, I. Kolosenko, Á. Végvári, Å.M. Wheelock

**Author notes:** Corresponding Authors: Martina Bonatti: Karolinska Institutet, CMM L8:02, Karolinska University Hospital Solna. 171 76 Stockholm, Sweden., Åsa M. Wheelock: Karolinska Institutet, CMM L8:02,Karolinska University Hospital Solna. 171 76 Stockholm, Sweden.

## Abstract

**Background:** Early detection and longitudinal follow-up are essential for timely diagnosis and treatment for lung diseases. Currently, invasive methods are often required to examine the distal parts of the lungs. The growing need to explore the bio-molecular mechanisms in various lung diseases highlights the importance of non-invasive methods. The use of particles in exhaled air (PExA), a non-invasive technique for sampling of epithelial lining fluid from distal airways, is gaining attention. MicroRNAs (miRNAs) are crucial in modulating protein expression both intracellularly and intercellularly, often transported via extracellular vesicles. Dysregulated miRNAs have been linked to many pulmonary diseases, and their relative stability, especially when encapsulated in EVs, makes them promising biomarkers. Here we report for the first time multi-modal analysis of miRNAs and proteins in PExA, offering an opportunity to study the role of miRNAs in the pathophysiology of respiratory diseases in a non-invasive manner.

**Methods:** Exhaled particles were collected from healthy subjects using the PExA 2.0 instrument utilizing the optimized PExA breathing maneuver. PExA samples collected on different types of impaction membranes were analyzed using a non-targeted mass spectrometry-based proteomics workflow optimized for single-cell detection, and a miRNAseq workflow optimized for low input starting material. Technical validation of a subset of the detected miRNAs was performed using custom-designed miRCURY LNA miRNA PCR assays. Pathway enrichment analyses for the detected proteins were performed using STRING.

**Results:** Proteomic analysis consistently identified over 50 proteins across multiple types of impaction membranes, sample dilution series, and individuals down to a single PEx sPOT (24ng starting material). We observed a significant enrichment of proteins associated with extracellular vesicles, including blood microparticles, and secretory granules. miRNA-seq revealed 39 mature miRNAs, the majority of which have been previously reported to be detected in the airways. Some were also reported to be secreted by primary human airway epithelial cells via extracellular vesicles. miRNA-125b and the members of the let-7 family were among the most abundant miRNAs detected. Fluorometric assays showed significant RNase activity in both PExA and other lung-related samples, such as bronchoalveolar lavage fluid, suggesting that this activity originates from the airways and is independent of the sampling techniques used. The workflow for extraction and processing of the PExA collection membrane, tested with abundant synthetic miRNAs and analyzed using the miRCURY LNA miRNA PCR assay, yielded results comparable to control samples, indicating that the membrane material does not interfere with the assay.

**Conclusions:** Using PExA, we identified several miRNAs reported to be dysregulated in pulmonary disorders. The enrichment of extracellular secretory components in the core protein list, along with the elevated RNAse activity in the respiratory tract, suggest that the detected miRNAs may be encapsulated within extracellular vesicles. These miRNAs are of particular interest due to their potential role in intercellular communication. Our findings suggest that PExA holds a potential as a non-invasive tool for studying extracellular vesicle-mediated miRNA cargo in the small airways.

## INTRODUCTION

Early detection of chronic inflammatory lung diseases is key to avoid loss of lung function and to maintain quality of life for affected individuals. In the case of lung cancer the need for early detection is even more pronounced, with detection prior to first symptoms being highly associated with survival. Today, we rely on invasive sampling techniques such as lung biopsies or bronchoalveolar lavage (BAL) for diagnosis and biomarker discovery. The invasiveness makes sampling resource intensive and impractical for longitudinal studies; also, bronchoscopy-mediated sampling may not be well-tolerated for some patient groups. It is thus essential to balance the benefits with the potential risks and patient discomfort associated with these procedures (1). As such, the development of non-invasive sampling techniques of the distal lung is pivotal to facilitate biomarker discovery, longitudinal monitoring or large-scale screening of respiratory disease. Particles in exhaled air (PExA) represents a unique sampling method that permits non-invasive sampling of the epithelial lining fluid and airway exudates from the small airways and alveoli (2). This sample type is also referred to as liquid biopsies from the distal airways. The approach involves impaction of exhaled aerosols comprising ultrafine droplets of respiratory tract lining fluid (RTLF) onto a membrane (3). These droplets are generated and exhaled after a specific breathing maneuver that facilitates the closure and reopening of the small airways. PEx samples have previously been shown to contain several types of biomolecules, including proteins (2, 4) and lipids (5) known to be present in the RTLF in the distal airways and alveolar region. Accordingly, this type of sampling method can be especially relevant for disorders involving the small airways, such as asthma and COPD (2, 6, 7). Despite several recent advancements of the sampling method and associated publications, our understanding of the material collected and its relevance to disease remains limited. As such, further benchmarking and analysis needs to unlock the full potential of this non-invasive technique.

MicroRNAs (miRNAs) are small, conserved non-coding RNAs that regulate gene expression post-transcriptionally (8). They play a crucial role in various biological processes, and their dysregulation is linked to many pulmonary diseases (9–11). miRNAs are found in numerous body fluids and extracellular spaces, including RTLF, either associated with other biomolecules or encapsulated in extracellular vesicles (EVs) (12–14). Acting as functional vehicles of intercellular communication, EVs transport miRNAs during pathological processes, protecting them from degradation, and thereby enhancing their stability (15). Given their importance in diseases processes (9, 10), as well as relative stability compared to both proteins and other RNA species, miRNAs hold great potential for various clinical applications, both as diagnostic and prognostic biomarkers and for longitudinal monitoring of disease progression and treatments. Non-invasive molecular tools for studying lung-derived miRNAs are emerging, though not yet rigorously evaluated or in clinical use (16). Existing publications on non-invasive approaches focus on exhaled miRNAs, primarily using exhaled breath condensate (EBC) for sampling (16–18). Radioactive markers studies revealed that EBC predominantly captures central airways, in contrast to the PExA sampling method which appears to sample primarily RLTF from the smaller airways (19).

Given the pivotal roles of extracellular vesicles (EVs) and miRNAs in intercellular communication and regulatory processes in lung diseases, we have developed a miRNA analysis approach for non-invasive sampling of the distal lung using the PExA technology. Here we present a benchmarking of the workflow in healthy subjects across multiple membrane matrices for PEx impaction, exploring the detection limit through a dilution of the PEx SPoT punch technology. In parallel, a robust sensitive non-targeted proteomics workflow originally developed for single-cell analyses was benchmarked to be used in conjunction.

## MATERIALS AND METHODS

### Sample collection

PEx biospecimen were collected using the PExA 2.0 instrument (PExA AB, Gothenburg, Sweden), using standardized methods adapted from previously described methods (20, 21). The collection was performed by impaction on dried blood spot (DBS) membranes (Whatman 903 Protein saver, specially tailored for PExA, PExA AB, Gothenburg, Sweden) compatible with the sPOTs punching technology, the original PExA teflon membranes, or Flinders Technology Associates (FTA) membranes (QIAcard FTA Elute Indicating Micro, Qiagen, specially tailored for PExA, PExA AB), a chemically coated cellulose membrane designed to lyse cells on contact, denature proteins and protect nucleic acids from degradation. All subjects performed a standardized breathing maneuver (22, 23) using a mouthpiece equipped with a two-way, non-rebreathing valve (Series 1420; Hans Rudolph, Shawnee, KS, USA), wearing a nasal clip during the process. Prior to the sampling, the subjects took a minimum of three tidal breaths of HEPA-filtered air via the PExA instrument to minimize particle contamination from surrounding ambient air. From each subject, 240 ng of exhaled particles were collected. Following the collection process, the sample holder containing the impaction membrane was directly placed in a dust- and RNAse-free container, and the area of the membrane impacted by the particles (PEx sPOT) were excised using a puncher specifically designed for this purpose. Subsequently, the PEx sPOTs were transferred to 1.5 ml Eppendorf® DNA LoBind tubes for miRNA analysis, or Protein LoBind tubes for proteomic analyses, and stored at -80°C until use. All the procedure to prepare the sample holder and to collect the PEx sPOTs were performed under a laminar flow hood under aseptic conditions to prevent sample contamination.

### Proteomics workflow

Two types of sample substrates were evaluated for PEx material collection for proteomics analyses: The teflone membrane developed for the original PExA instrument, made of polytetrafluoroethylene (PTFE), featuring a 0.45 μm pore size, 25 mm diameter (Merck Millipore Ltd., Cork, Ireland), as well as the substrate optimized for the sPOT punch technique made of cellulose, also known as the dried blood spot (DBS) membrane (Whatman 903 Protein saver, specially tailored for PExA). Sample punch-out technique (sPOT) was used for both substrates to excise membrane circles of equal sizes.

Two distinct PEx proteome experiments, outlined in Figure 1A, were designed to evaluate the technical reproducibility and the detection limit of protein identification in PEx material using both PTFE and DBS membranes. In Experiment 1, we collected 240 ng PEx sample from a healthy subject and divided it into three replicates composed of three PEx dots (i.e., 3 replicates of 72 ng PEx each). In Experiment 2, we collected the equivalent sample from the same subject on a different day, maintaining the same time of the day to minimize potential diurnal variation, dividing it into five different replicates, each composed of a single PEx sPOT (5 replicates of 24 ng PEx). Subsequently, proteome analyses were performed as described below.

**Figure 1.**
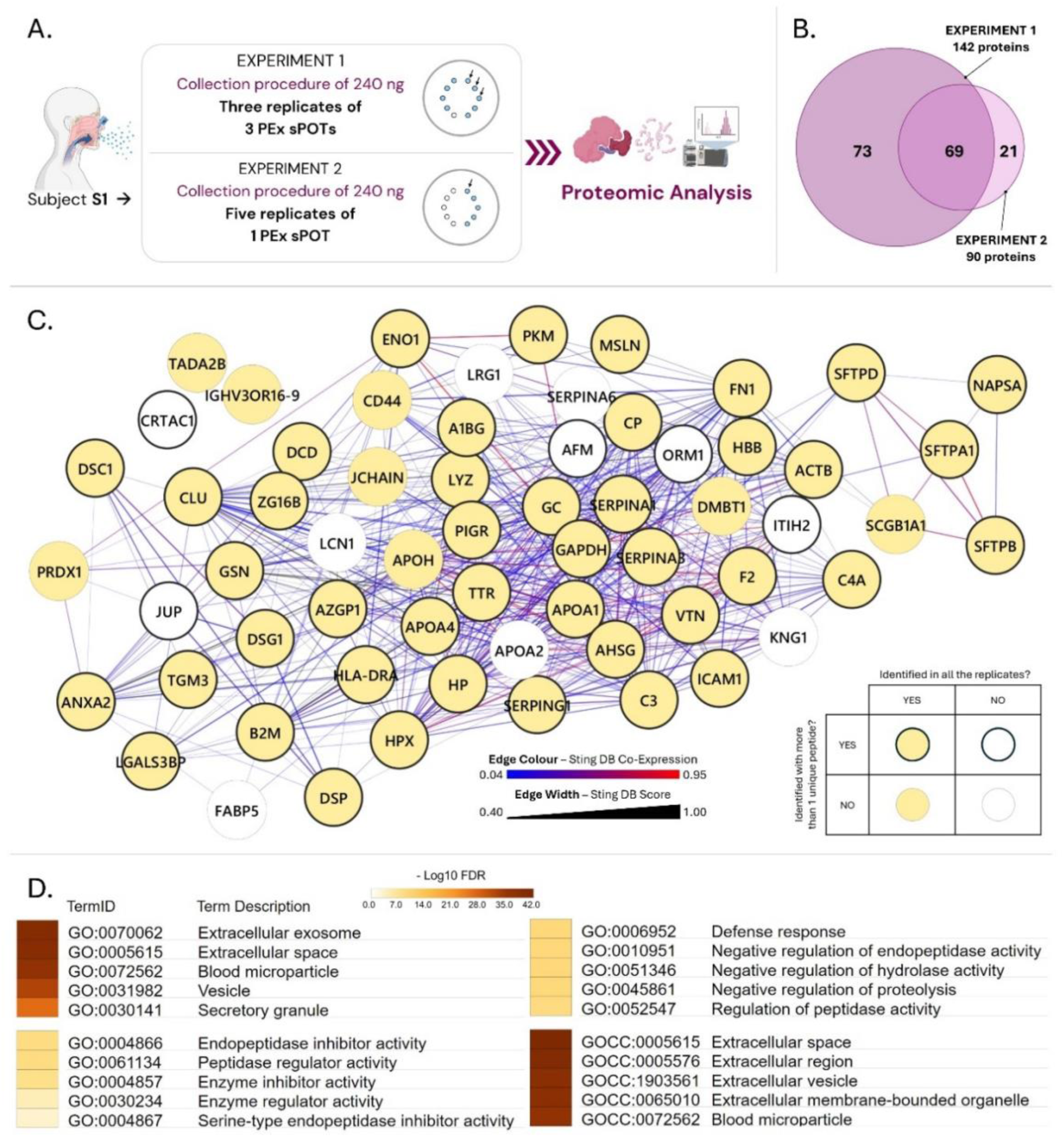
Proteins detected in PEx and their associated Gene Ontology (GO) categories. **A.** Graphical outline of the experimental workflow. **B.** Venn diagram displaying the number of proteins identified across all replicates in both experiments (Experiment 1: triplicate of 72 ng PEx starting material; Experiment 2: 5 replicates of 24 ng PEx starting material). **C.** The protein-protein interaction (PPI) network of proteins identified in all replicates of both experiments. The legend describes the features of the network’s elements. Edges represent the StringDB score, describing the confidence of the corresponding PPI. For the network realization, the minimum required interaction score was set to 0.4 (medium confidence). Edges are colored based on the co-expression interaction score (See legend below graph; Light grey: no interaction, red: high, blue: low), and thickness of edges represents String DB score. Yellow nodes with black border represent proteins with the highest level of confidence for identification (identified in all replicates with minimum 2 unique peptides/protein). **D.** Pathway enrichment analysis based of proteins identified in both experiments, with the top five enriched terms for all three GO categories and subcellular compartments displayed. Color codes display the significance scores (FDR in-Log10 scale). All data were obtained from stringDB (25).

### Non-targeted proteome analysis

Excised PEx membranes spots were wetted with a total volume of 20 µL lysis buffer composed of 0.2% n-dodecyl-β-D-maltoside (DDM; Sigma-Aldrich) in 100 mM triethylammonium bicarbonate (TEAB; Fluka) and sonicated in a water bath for 10 min. Suspensions were vortexed and centrifuged at low speed for 5 min at room temperature to sediment PEx sPOTs. Subsequently, samples were digested by adding 10 µL of 20 ng/µL sequencing grade trypsin (Promega) into the lysis buffer, with a final trypsin concentration of 6.7 ng/µL. Following a short vortexing samples were incubated for 4 h at 37°C and centrifuged at 10,000 x *g* for 5 min. The supernatants were transferred to sample vials and dried in a vacuum concentrator (Eppendorf).

Peptides were reconstituted in solvent A (0.1% formic acid, FA in 2% acetonitrile, ACN) and 2 µL of each sample was injected using an Ultimate 3000 nano-flow UHPLC system (Thermo Fisher Scientific). Peptides were captured on a 2 cm Acclaim PepMap trap column (Thermo Fisher Scientific) and separated on a heated 25 cm long EASY-Spray C18 column (Thermo Fisher Scientific) applying a 90 min long gradient: 4-26% of solvent B (0.1% FA in 98% ACN) in 90 min, 26-95% in 5 min, and 95% of solvent B for 5 min at a flow rate of 300 nL/min.

Mass spectra were acquired on an Orbitrap Eclipse tribrid mass spectrometer (Thermo Fisher Scientific) equipped with high-field asymmetric ion mobility spectrometry (FAIMS) in *m/z* 300 to 1500 at resolution of R=120,000 (at *m/z* 200) for full mass, targeting 8×10^5^ ions in maximum 100 ms, followed by data-dependent higher-energy collisional dissociation (HCD) fragmentations of precursor ions with a charge state 2+ to 6+, using 20 s dynamic exclusion. The tandem mass spectra of the top precursor ions were acquired in 1 s cycle time by the linear ion trap in rapid mode, targeting 3×10^4^ ions for maximum injection time of 20 ms, setting quadrupole isolation width to 1 Th and normalized collision energy (HCD) to 30%. FAIMS was operated at three different compensation voltages in each cycle using either -40, -60 or -80 V.

### Proteome data analysis

The raw files were searched against the human UniProt database using MS Amanda v2.0 search engine (24) loaded into Proteome Discoverer 2.5 software (Thermo Fisher Scientific). MS1 precursor mass tolerance was set at 10 ppm, and MS2 tolerance was set at 0.6 Da. Search criteria included variable modifications of oxidation (+15.9949 Da) on methionine residues, deamidation (+0.984 Da) asparagine and glutamine residues. Search was performed with full trypsin/P digestion and allowed a maximum of two missed cleavages on the peptides analyzed from the sequence database. The false-discovery rates of proteins and peptides were set at 0.05. All data were quantified using the label-free quantitation node of Precursor Ions Quantifier through the Proteome Discoverer calculating protein abundances based on precursor abundances. The abundance levels of all proteins were normalized using the total peptide amount normalization node in the Proteome Discoverer.

The technical reproducibility of the proteomics analyses across replicates was assessed for proteins identified as Master proteins, and detected across all five technical replicates for each membrane type. Protein abundances were normalized by a sample-specific normalization factor calculated as the [sample total protein abundance]s/[median total protein abundance across samples], calculated separately for each membrane type. Normalized abundances were log2-transformed for downstream analyses where indicated.

### Protein-protein interaction (PPI) Network and Pathway analysis

Pathway enrichment analysis and PPI network visualization was performed using STRING database v. 12.0 (25). The 69 proteins identified reproducibly across Experiments 1 and 2 were included in the analysis. Keratins were eliminated due to possible contamination concerns, and a few protein IDs were not found in the STRING database, resulting in the final pathway analysis and visualization network being based on 62 proteins. For the PPI network, the minimum required interaction score was set to medium confidence (cutoff 0.4). The obtained network was then transferred to Cytoscape v. 3.9.1 (26) for graphical processing.

### miRNA workflow

Two experiments were designed to evaluate miRNA expression in PEx material using DBS membrane. Two independent PExA samples of 240 ng each were collected from two healthy subjects, and PEx sPOTs were excised. RNA extraction was performed from material from 10, 2 and 1 sPOTs respectively for each subject (see Figure 2A) as described below to determine the detection limit for miRNA quantification of PExA samples.

**Figure 2.**
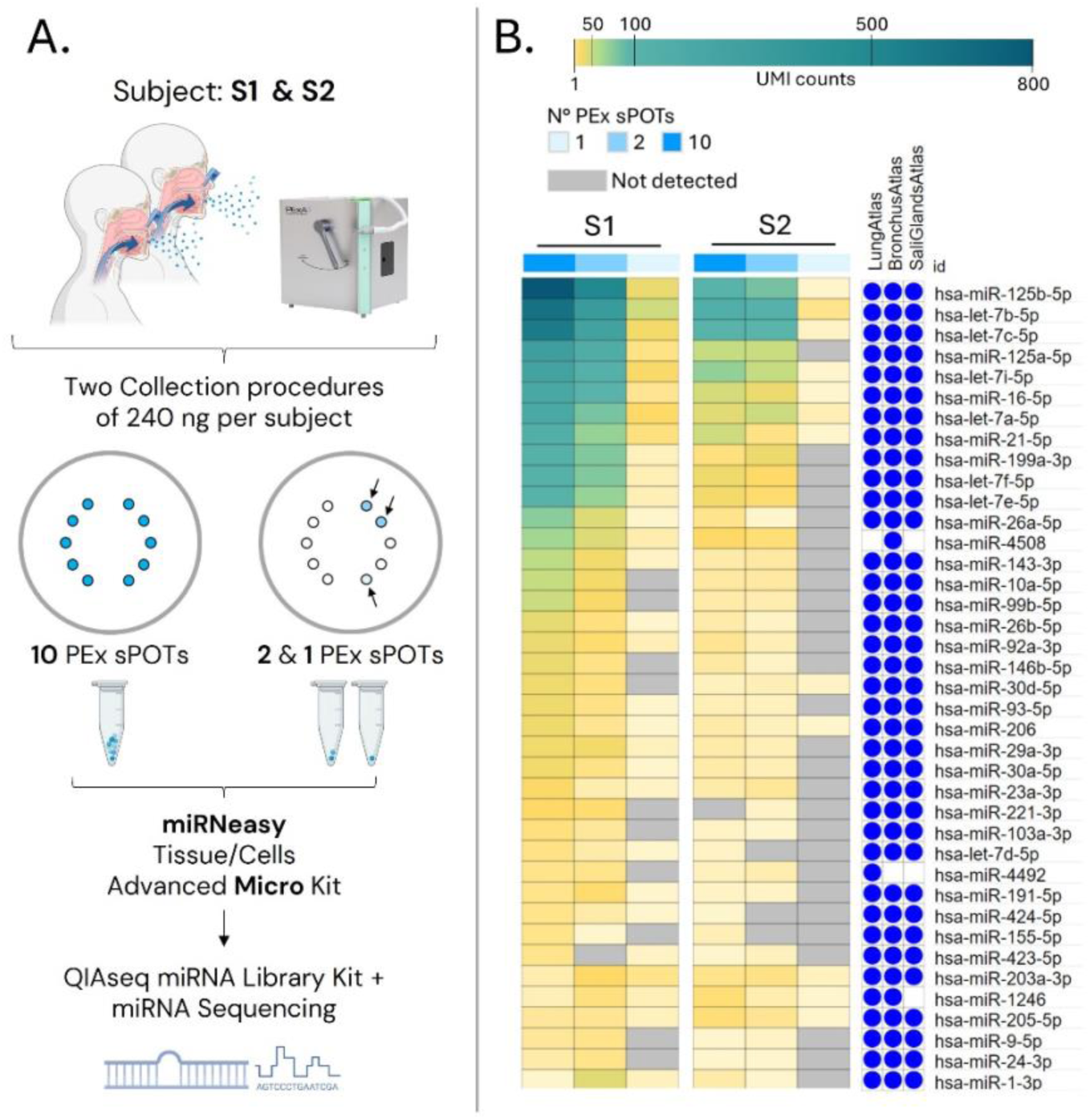
miRNA identified in PEx samples on DBS membrane using RNAseq. **A.** Graphical outline of the experimental workflow. **B.** List of detected miRNAs. Data are presented as UMI counts color, coded from dark green (UMI=800) to light yellow (UMI=1), with light green representing UMI>50. Each of the two subjects are represented by 3 columns, related to 10, 2 and 1 PEx sPOTs of starting material respectively. The right columns display gene lists sourced from or miRNATissueAtlas2 (with expression mean > 10 as cut-off (27, 28)). A blue dot indicates the presence of the PExA-identified miRNA in the corresponding database.

### RNA extraction

Vials containing PEx sPOT were transferred directly from -80°C storage to dry ice to prevent premature thawing, and 260 μL of Buffer RLT (Qiagen) was added while still in a frozen state. Samples were transferred to room temperature and incubated for 10 min with intermittent vortexing, then mixed through pipetting until PEx sPOT were disintegrated, thereby ensuring the creation of a pseudo-uniform solution. Total RNA, including the miRNA fraction, was extracted using the miRNeasy Tissue/Cells Advanced Micro Kit (Qiagen) according to manufacturer’s instruction. The gDNA Eliminator Spin Column step is essential as it allows to separate any paper-based particles from the liquid, that could interfere with any downstream steps. The RNA fraction was eluted in 15 µL nuclease-free water, and the process of elution was performed twice with an incubation time of 5 min each to increase the elution efficiency. For samples destined for the qRT-PCR-based miRNA assays, 1µl of RNA spike-in mix (Qiagen) was added to the lysis buffer during RNA isolation to monitor RNA extraction efficacy.

### Library Preparation and RNA sequencing

The QIAseq miRNA Library Kit (Qiagen, Hilden, Germany) was used to prepare the library, which has integrated Unique Molecular Indices (UMIs) to facilitate measurement of pre-amplification miRNA levels. The libraries were sequenced on the NextSeq 2000 instrument (Illumina, San Diego, US) with a 100 nt (Read1) 8 nt (Index1) setup using ’P2’ flowcell, at an average sequencing depth of 26 M reads/sample. The fastq data were uploaded to the RNA-seq Analysis Portal (RAP) 5.0 powered by QIAGEN CLC Genomics Workbench to obtain the UMI raw counts (reference: miRBase_v22, Homo sapiens; analysis workflow version: 1.2).

### cDNA Synthesis and qRT-PCR based miRNA assays

For validation experiments using qRT-PCR based miRNA assays, a set of samples were collected in an identical manner as for the RNAseq analysis described above. Following extraction, RNA was reverse transcribed using miRCURY Locked Nucleic Acid (LNA™) Reverse Transcription (RT) kit (Qiagen) according to the manufacturer’s instruction. For all samples, the reaction mix included the maximum allowable amount of RNA (6.5µL) in a final volume of 10 µL. The cDNA synthesis control (UniSp6) and the PCR amplification Ctrl (cel-miR-39-3p, called UniSp3) coming from the Spike-in mix (Qiagen) were introduced to the reverse transcription reaction to assess its efficacy. A miRCURY LNA miRNA Custom PCR Panel (Qiagen) was used and the set of primers for the miRNAs selected for the custom plate were all validated for qPCR. cDNA samples were amplified using miRCURY LNA™ SYBR Green PCR kit (Qiagen) and the real-time reaction was performed using a QuantStudio™ 7 Flex Real-Time PCR System (Applied Biosystem Foster City, CA, USA), with the following protocol: 95°C for 2 min, followed by 40 cycles consisting of denaturation (10 s at 95°C) and combined annealing/extension (60 s at 56°C). Melting curve analysis from 60–95°C was performed post-amplification to evaluate product specificity.

### RNAse activity assay

The presence of RNase activity in bronchoalveolar lavage fluid (BALF) and PEx samples was assayed using the RNaseAlert® Lab Test Kit (Thermo Fisher Scientific). BALF samples were kept undiluted and thawed on ice when frozen. The assays were conducted following the manufacturer’s protocol, with fluorescence measured using the Varioskan™ LUX multimode microplate reader at an excitation wavelength of 490 nm. As an additional control reference sample, DBS membranes were pre-treated with various RNAse inactivating reagents prior to sample collection, including RNaseZap™ (ThermoFisher Scientific) and lysis buffer from the miRNeasy Tissue/Cells Advanced Micro kit (Qiagen).

### Evaluation of the potential interference of the membrane in the extraction of the RNA

To determine if the membrane matrix materials interfere with the RNA extraction procedure and workflow, 5 PEx sPOTs each of DBS and FTA membrane were wetted with 0.2 µL of RNA spike-in mix (Qiagen), making a total of 1 µL across 5 punches. RNA extraction was carried out as described above. To control for adsorption effects, a positive control was performed by adding 1 µL of the RNA spike-in mix directly to lysis buffer, without including any PEx sPOTs membrane.

### Ethical Statement

Analysis of miRNA and proteins collected by the PExA method has been approved by the Swedish Ethical Authorities (Case no. 2022-03768-01). For the purpose of methods optimizations, PEx material was stripped of all personal identifiers at the time of collection, and all downstream handling and analysis was performed in a de-identified manner.

## RESULTS

### Proteomics analysis of PExA material

The quantitative performance of a non-targeted, mass-spectrometry based proteomics of PEx material was assessed across substrate types and material scarcity based one i) number of detectible proteins and ii) quantitative variance across technical replicates. Specifically, the performance of material collected on PTFE vs DBS substrates using 3 vs 1 PEx sPOTs of starting material were assessed (See Figure 1A: 3 replicates of 72 ng vs 5 replicates and 24 ng). A third membrane type optimized for RNA sampling was also evaluated, but yielded no protein. The technical reproducibility was comparable for the two membrane types both in terms of normalized abundance (**Figure S1A-B, S1E-F**), and CVs (**Figure S1C-D, S1G-H**). Importantly, the correlation between log2-normalized protein abundances and CVs was comparable between membrane types with the exception of a handful of proteins with a somewhat higher CV for the DBS membrane (Pearson r > 0.90). Overall the technical replication supports a high level of reproducibility of the proteomic measurements even when amounts as scarce as a single PEx spot were analyzed (**Figure S1G-H**). Also the number of proteins identified was comparable across membrane types, with high overlap between the replicates (**Figure S2A-B**). Overall the performance of the DBS membrane, a newer PExA collection substrate optimized for the sPOT punching technique, was compared to that of the original, more widely used PTFE membrane, demonstrated comparable protein abundance distributions, inter-replicate correlations, and technical variability. Considering the practical advantages of using DBS, it may be used as a convenient alternative to PTFE membranes in proteomics workflows.

Non-targeted proteome analysis of all replicates of the two experiments revealed a total of 163 proteins detectable in DBS PExA membranes, with 69 proteins in common between the two experiments (**Figure 1A**). We identified 142 proteins in the first experiment (72 with more than 1 unique peptide) and 90 in the second (44 with more than 1 unique peptide). Of the 69 common proteins, 58 were identified in all replicates of both experiments. All the proteins in this list that are present in the STRING database are shown in **Figure 1B**, relevant information and detailed gene names are reported in supplementary material (**Table S1**). The provided list showed statistically significant enrichment (false discovery rate, FDR ≤ 0.05) for several terms associated with extracellular secretion-related components, including extracellular exosome, blood micro-particle, vesicle and secretory granule (**Figure 1C**). A complete list and further details of the pathway analysis are reported in the supplementary materials (**Table S2**). On one PEx sPOTs, 70 proteins were identified in all the 5 replicates, of which 43 were identified with more than one unique peptide, that gave statistically significant enrichment for terms associated with extracellular release processes (data showed in supplementary material, **Table S3**), providing a confidence of the analysis performed even in one single PEx sPOT, here representing a starting material of 24 ng PEx.

### miRNA content evaluation on DBS membrane using miRNA NGS sequencing

Subsequently, we sequenced the PEx material to analyze the miRNA composition. A total of 181 miRNAs were detected with at least 1 UMI count in at least 1 PEx sample. A set of 39 miRNAs were detected with at least 10 UMI counts in at least one of the samples (**Figure 2B**). As expected, the UMI counts are higher in the samples containing material from 10 PEx sPOTs and then they gradually decrease in the samples processed with 2 and 1 PEx sPOTs for both subjects. The number of counts in the sample with 1 PEx sPOTs are low or absent for most of the miRNAs. Most miRNAs detected in PEx material are known to be present in respiratory tract miRNA atlases (lungs, bronchus, and saliva (27, 28)), **Figure 2B, right panel.** The highest UMI counts were registered for miR-125b-5p in the 10 PEx sPOTs of subject 1, and then the number gradually decreased in the 2 and 1 PEx sPOT. In subject 2 we observed the same trend, but with lower counts. We detected several members of the let-7 miRNA family (let-7a-5p, let-7b-5p, let-7c-5p, let-7e-5p, let-7f-5p, let-7i-5p) with high UMI counts. They exhibit the same reduction trend as miR-125b-5p, transitioning from 10 to 2 to 1 PEx sPOTs.

### RNA degradation evaluation in lung related samples

During the QC analysis of miRNA-seq data (data not shown), a significant proportion of reads that were shorter than expected mapped to RNAs other than miRNA, indicating potential RNA degradation. We then performed a fluorimetric assay to evaluate the presence of RNAse activity in the airways to exclude any contamination coming from the PExA instrumentation, tools or workflow. RNase activity was measured in PExA samples collected on a DBS membrane, with fresh and frozen BALF samples serving as a baseline control for inherent RNAse activity originating from the airways compartment (**Figure 3A**). All samples, both PEx and BALF, displayed significantly higher fluorescence signal than the negative control, indicating strong RNase activity. We then measured the RNAse activity on the membrane only and applying some RNAse inactivation strategies (**Figure 3B**). The fluorescence level of the DBS membrane alone was comparable to the negative control. Samples collected on DBS without any protective reagents resulted in being contaminated by RNAses, even if the activity was a less in the samples that were immediately frozen at -80°C after the collection. In samples that were collected on DBS that had been pre-treated with lysis buffer from the RNA extraction kit, or in samples that were collected on an FTA membrane the RNAse contamination is minimal and comparable to the negative control.

**Figure 3.**
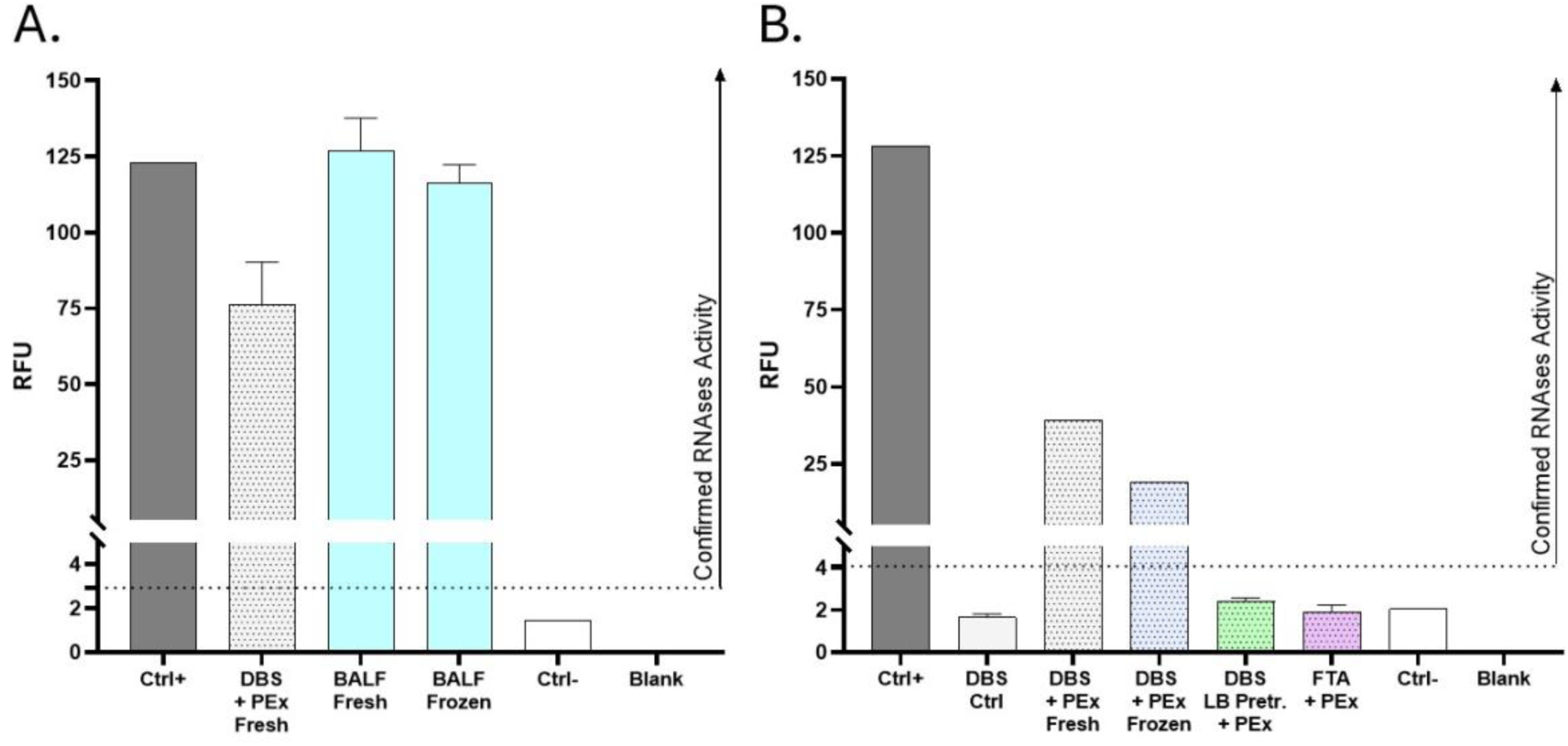
Evaluation of RNAse activity in samples collected from the airways by means of PExA and BALF. **A.** Bar graph showing the level of RNAse activity in different sample types collected from the airways, including PEx samples collected on DBS membrane, fresh BALF, and frozen BALF. **B.** Bar graph showing the level of RNAse activity in PEx samples using different condition and membrane matrices (Ctrl +: positive control provided by the manufacturer of the kit, with known concentration of RNAse; DBS Ctrl: DBS membrane without sample; LB Pretr. = DBS membrane pretreated with lysis buffer from the RNA extraction kit; Ctrl -: negative control, reaction containing RNase-free water). The dotted line represents fluorescence levels exceeding two-fold the corresponding negative control, defined as the threshold for detected RNase activity. Data are reported as mean + SD. The dotted texture highlights samples with PEx material.

### Validation of miRNA using real-time PCR technology

A subset of the miRNAs identified with RNA-seq were validated with a targeted qRT-PCR based miRNA assays, with CT values less than 35 (**Figure 4A**). MiR-125b-5p appears to be the most abundant based on the clear melting curve plot (**Figure 4B**), as reported in the RNAseq experiment. However, some miRNAs have not been identified using this technique. The internal controls for RNA extraction (UniSp2 and UniSp4), cDNA synthesis (UniSp6), and PCR amplification (UniSp3) appear to be reliable and uniform across the different samples, ensuring a high level of reproducibility in the overall workflow (**Table S4**).

**Figure 4.**
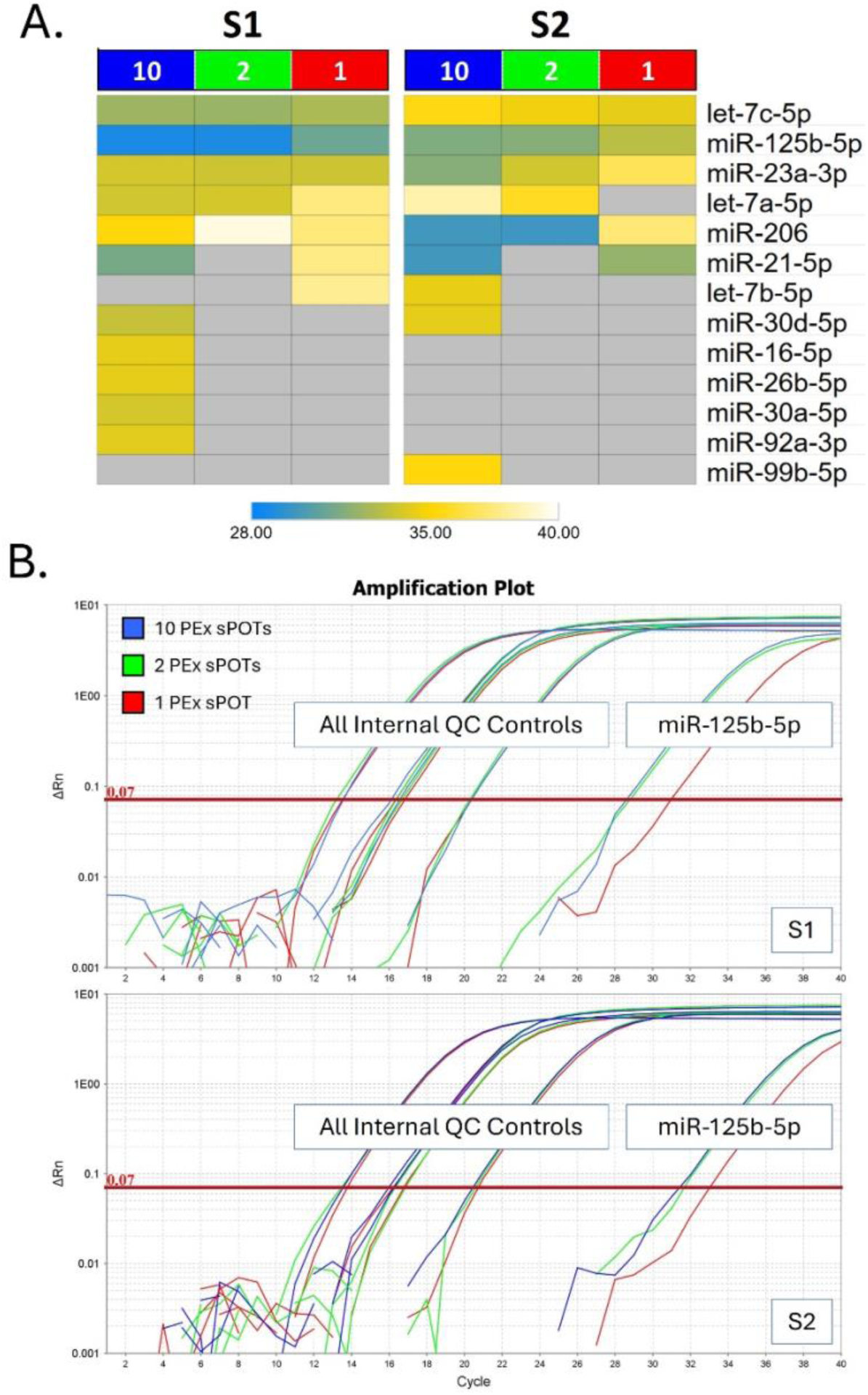
Technical validation of miRNA detection in PEx samples using a custom designed miRCURY LNA™ SYBR Green PCR assay. Results of the PCR array performed on the same set of samples utilized in the RNAseq analysis. In spite of the inferior detection limit of the PCR assay compared to the UMI-corrected miRNA-seq methodology displayed in Figure 2, the presence and reproducibility across subjects could be confirmed for several miRNAs even at the lowest amount tested (24 ng PEx material). **A.** Visualization of miRNA presence across a dilution series of 10, 2 and 1 sPOT (240 ng, 48ng and 24 ng PEx material) from 2 subjects. Grey cells represent miRNA below detection limit for the assay, and colors represent number of cycles required. **B.** Amplification plot of the internal controls of the array as well as of the most abundant miRNA in S1 and S2 (miR125b-5p).

### Effect of PExA membranes on RNA extraction

To evaluate if the DBS or FTA membrane matrixes interfere with the workflow, we used the spike-in mix used during the main experiment. Given the extremely scarce amount of RNA present in PEx samples, the spike-in mix from the qRT-PCR added directly to pre-cut PEx sPOTs rather than standard PEx material was used to evaluate variability of the collection procedures using the miRCURY LNA miRNA PCR assay. CT values for the five synthetic miRNAs of the spike-in kit (UniSp5, UniSp4, UniSp2 as RNA extraction controls, UniSp6 for cDNA synthesis, and UniSp3 for RT-PCR) were comparable to controls where the mix was processed without membranes (**Figure 5**). Thus, the DBS and FTA membranes do not interfere with RNA extraction nor affect the enzymes used in cDNA synthesis and RT-PCR.

**Figure 5.**
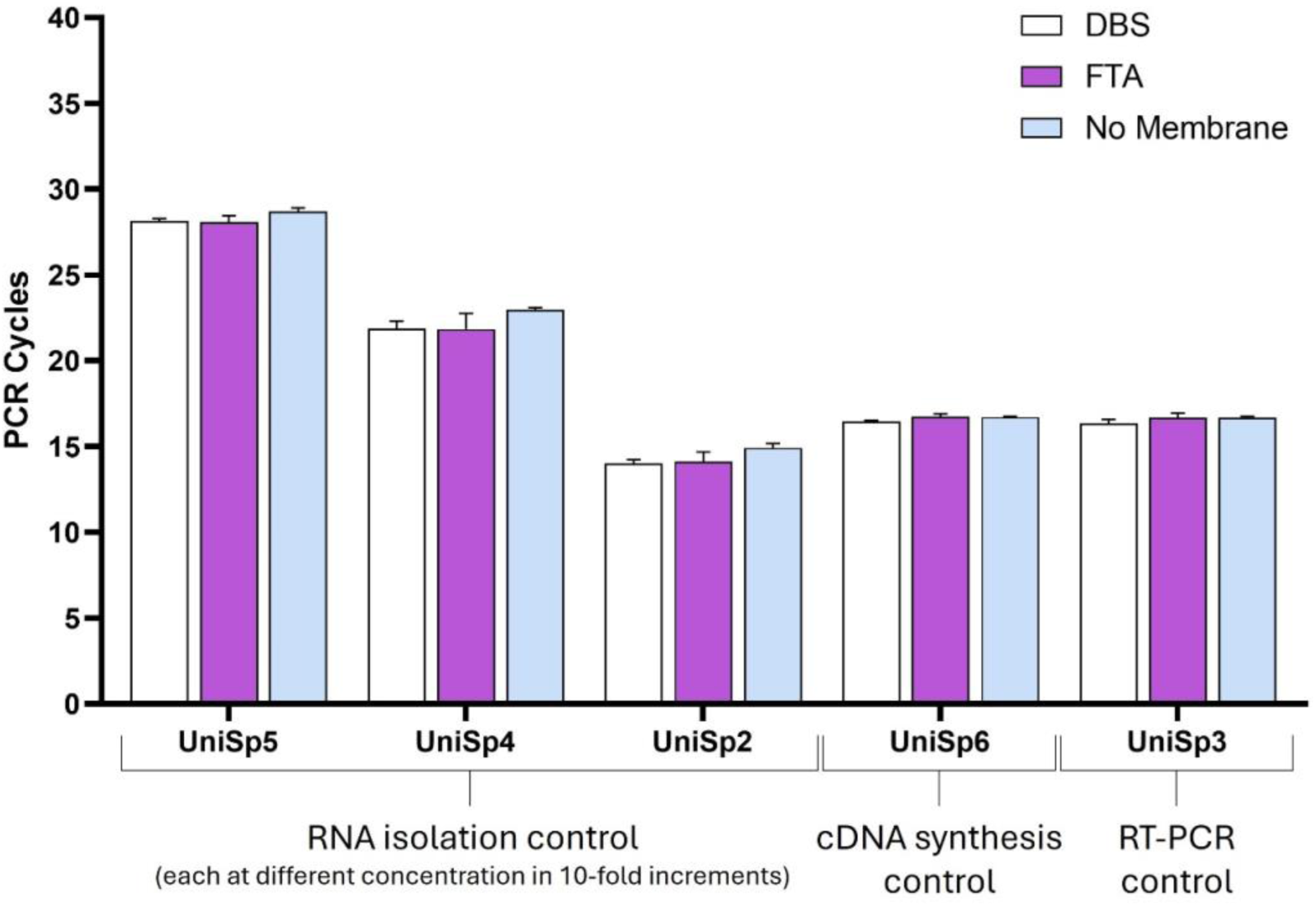
Evaluation of potential membrane interference during RNA extraction. PCR cycle thresholds for RNA spike-in controls (UniSp5, UniSp4, UniSp2) for RNA isolation, cDNA synthesis (UniSp6), and RT-PCR (UniSp3) show comparable results between DBS (white), FTA (purple), and no membrane (grey) conditions, with no statistically significant differences between conditions (Kruskal-Wallis test). Data are reported as mean + SD across three replicates.

## DISCUSSION

These studies were designed to evaluate the feasibility of multi-omics analysis of PExA samples, with the focus on establishing a workflow for MS-based proteomics and miRNA-seq from a single sample. We report for the first time the presence of miRNAs in material collected using the PExA methodology, and provide novel insights into the existence of proteins and miRNAs of EV origin in these non-invasive samples from the distal airways. A total of 181 miRNAs were detected in PEx sPOTs using RNA-seq analysis, including 39 miRNAs identified with high confidence, and a subset further verified with qPCR. In parallel, 163 proteins were detected using non-targeted proteomics, with 69 proteins identified reproducibly across subjects with scarce amounts of starting material. Pathway enrichment analyses revealed that a large proportion of these proteins are associated with extracellular vesicles and secretory pathways, indicating the presence of EVs. Moreover, RNase activity measurements in PEx samples as well as BALF samples indicating that the sequenced miRNAs are of intravesicular origin.

PExA has recently gained increasing attention as a non-invasive method for collecting material from the distal lung compartment to study molecular changes in health and disease. Commonly used sampling techniques for collection of biomolecules from the lung, such as bronchoscopy and induced sputum, are inherently invasive sampling methods that require extensive clinical and pre-clinical logistics, as well as exert discomfort for research subjects. In addition, their invasive nature makes repeated sampling and longitudinal follow-up challenging. The majority of less invasive sampling techniques available in respiratory medicine, such as EBC and volatile organic compound (VOC) in exhaled breath have been shown to primarily sample from the proximal airways. In contrast, PEx particles have been shown to originate from the RTLF of the distal airway and alveolar region (2, 4) with the strongest evidence coming from the detection of surfactant proteins predominantly produced in the distal airways (21, 29). It has also been shown that the protein content of PEx particles is rather stable, making them promising for identification and monitoring of protein biomarkers of lung pathologies (4).

A number of previous publications (2, 4, 19–21) have reported the presence of proteins in PEx samples. In this study, we performed non-targeted proteome analyses with the aim of investigating the minimal quantity of PEx material required for analysis without compromising the reproducibility of the assay. Our analyses of the PEx proteome demonstrated a significant enrichment of GO terms related to EVs, including exosomes. Our findings thus further strengthen the notion that the protein content of PEx can be identified with mass spectrometry-based methods and that PEx samples contain EVs, including EVs of exosomal origin (2). Furthermore, we showed that this information can be captured even when analyzing a single PEx sPOT containing 24 ng material. Given that 10 PEx sPOTs that can be retrieved from a PEx membrane, this observation paves the way for the analysis of multiple classes of biomolecules (e.g., proteins, lipids, RNAs) from a single sampling occasion, which previously have required several sample sessions from each subject. Furthermore, our proteomics analyses exploring different PExA membrane matrices verified that the use of PTFE membranes used in the classical PExA method, as well as the DBS membranes compatible with the sPOT punching technology both yield similar results in MS-based proteome analyses. In contrast, FTA membranes optimized for RNA and DNA analyses were not compatible with proteome analysis, yielding zero proteins (data not shown).

It has previously been reported that EVs can function as a delivery system for miRNAs and facilitate intercellular communication in the respiratory tract (9, 13), and numerous fundamental physiological processes in health and diseases are regulated by miRNAs (7–9). Our group as well as others have detected exosomes and their miRNA cargo in various lung samples, including BALF and sputum supernatants in patients with asthma and COPD, as well as in healthy subjects (14, 30). Cells are known to adjust the production and composition of EVs in response to various stimuli, and quantitative analysis of their cargo may thus provide useful insights into pulmonary disorders (30). The observation that a significant portion of the PEx proteome is associated with EVs suggests that intact miRNAs could be present in such samples, as their miRNA cargo would be protected from degradation (11, 31). Shi et al. provided a comprehensive report on exhaled miRNAs in lung cancer research utilizing the EBC sampling techniques (16). Specifically, they examine the efficacy of a selected panel of 24 miRNAs in differentiating individuals with non-small cell lung cancer from control subjects. Although further refinements of their exhaled miRNA interrogation technique are required, the researchers have shown that the detection of miRNAs in EBC using qPCR can lead to a modest enhancement in differentiating between cases and controls, beyond what can be achieved with clinical factor models alone (16). Nevertheless, the composition of biomolecules in EBC is significantly affected by their physical characteristics as well as their relative concentrations in the RTLF (32). Moreover, dilution poses uncertainty in EBC, as well as in BAL, where external fluid addition complicates concentration calculations and normalization. Conversely, PEx samples represent undiluted collection of droplets and particles from the RTLF of the distal airways and alveolar region, thus enhancing the reliability of quantification (4). For the above mentioned reasons, we explored the feasibility of performing miRNA-seq analysis of PEx samples in an attempt to performing multi-omics analyses of miRNA and proteomes from the distal airways in a non-invasive manner, and optimize its application for studying lung diseases.

The miRNAs identified through RNA-seq in PEx sample have previously been reported in multiple studies investigating miRNA profiles of the respiratory tract, including samples from lungs and bronchi reported in the Lung Atlas project (27, 28). In particular, the highly conserved let-7 miRNA family was detected in relatively high counts in both subjects investigated here, with let-7b-5p and let-7c-5p being the most abundant (33, 34). The significant level of preservation observed in let-7 family members across several animal species implies that they likely have crucial and maybe comparable functions in the biological processes of different organisms (33, 34). As summarized by Lee et al. (35), it appears that they exert two primary biological functions: crucial regulator of terminal differentiation and fundamental tumour suppressor. Furthermore, alterations of several let-7 family members in different respiratory diseases, including chronic obstructive pulmonary disease, idiopathic pulmonary fibrosis, and asthma (36) suggest their utility as biomarkers of respiratory disease. For example, based on the miRNA profile of exosomes derived from BALF, let-7 family members were included in a subset of miRNAs that distinguishes patients with asthma from healthy participants at baseline (30). The most common let-7 member found in PEx, let-7b-5p, was extensively studied by Koeppen et al. who demonstrated that EV-derived let-7b-5p reduced biofilm formation and increased antibiotic sensitivity in *P. aeruginosa* (36). Moreover, PEx samples contained four of the five most common miRNAs detected in airway epithelial cells-EVs by Koeppen and colleagues (36), which account for more than half of all miRNA sequence reads. Identification of miRNAs that are directly produced and encapsulated by airway epithelial cells provides further support for the concept that PEx samples can yield insights into intra-vesicular communication within the small airways. Other abundant miRNAs detected in PEx samples from both patients included miR-125b-5p and miR-125a-5p. The miR-125 family is found to regulate a set of genes that mediate cellular responses against pathogens through modulating, for example, IL-6 and type I IFN pathways (37). In addition, the miR-125 family has been shown to be implicated in lung carcinogenesis; however, the reports seem to find both oncogenic and tumor-suppressive functions of these miRNAs (37).

To address the potential problem of saliva contamination of the PEx samples, an issue that has been raised in the past for PExA, we compared the PEx miRNA content with that of the salivary glands (4). Three of the miRNAs detected in PEx, miR-4508, miR-1246, and miR-4492, are expressed in the bronchus and/or lung but not in the salivary glands based on the miRNA tissue atlas. Based on miRNA expression alone, we cannot completely exclude the possibility that our identified miRNAs originate from saliva. However, given that our PEx proteomic data did not detect amylase, the most prevalent protein in saliva often utilized as evidence to refute the presence of saliva, implies that no saliva was present in the PEx samples (4).

Given the frequent occurrence of RNA degradation in biological matrices (11, 31), we evaluated the presence of RNAse activity in PEx membranes following sample collection, as well as in BALF samples. The analyses confirmed high RNase activity. These findings confirm that the airway microenvironment is rich in RNases, suggesting that the miRNAs detected in PEx samples are indeed derived from EVs, which are known to protected their cargo from RNase activity and associated rapid degradation.

The presence of a selection of miRNAs detected by miRNA sequencing was validated using custom-designed miRCURY LNA™ SYBR Green PCR assay. Despite the low abundance of PEx material approaching the assay’s detection limit, the most highly abundant miRNA species, including miR-125b-5p and certain let-7 family members relevant to lung function, were detectable even in a single PEx spot also after UMI correction. Furthermore, our dilution series experiments showed that increasing the number of PEx sPOTs used for RNA extraction resulted in an increased miRNA UMI count for both individuals tested (Figure 2), therefore the use of 2-5 PEx SPOTs combined with a high sequencing depth is recommended. While several mature miRNAs were detectable in a single PEx sPOT, 2 or more PEx spots considerably increased the number of different miRNA species detected, as well as the qualitative information of the sampling. However, the dilution series experiments indicated that using the full panel of 10 sPOTs produced in a single sampling session may not be beneficial. Given these findings, we propose that designating five PEx sPOTs for protein analysis and five for miRNA analysis represents the optimal method moving forward. This ensures adequate material amount from an individual patient in a single PExA sampling session, facilitating the integration of protein and miRNA insights. Moreover, protein content may function as a prospective normalizing factor for miRNA data in the future, especially as additional research is required to identify universally stable miRNAs appropriate for quantitative normalization.

## CONCLUSION

This proof-of-concept study demonstrates that miRNAs can be detected in PEx samples and that the miRNAs identified can be relevant to use in studying and monitoring different lung diseases and associated treatment regimens. The enrichment of proteins from the EV- and exosome compartments along with the high level of RNAse activity detected both in BALF and PExA samples suggest that the detected miRNAs are derived from intact vesicles, which are known to offer protection via encapsulation. Taken together, we demonstrate the feasibility of the PExA method as a tool for the collection of non-invasive liquid biopsies to study EV-derived miRNAs from the small airways.

## Notes

### Competing Interest Statement

AMW is a Scientific Advisor for the company PExA AB. No research funding has been received from the company.

### Summary of Updates

General restiling for a better structure of the data for submission to specific journal

